# Wood structure explained by complex spatial source-sink interactions

**DOI:** 10.1101/2020.05.02.073643

**Authors:** Andrew D. Friend

## Abstract

Wood is a remarkable material. It is responsible for the sequestration of significant anthropogenic CO_2_^1^, aids understanding of past climates^2^, has unique acoustic, thermal, and strength properties, and is an endlessly renewable source of energy^3^. However, we lack a general integrated understanding of how its structure is created. A theoretical framework of wood formation is presented here that explains a diverse range of poorly understood observations, including: (i) the anatomy of growth rings, with a transition from low-density earlywood to high-density latewood; (ii) the high sensitivity of latewood density to temperature; (iii) cell-size regulation; and (iv) relationships between growth and temperature. These features arise from interactions in time and space between the production of cells, the dynamics of developmental zones, and the supply of carbohydrates. Carbohydrate distribution is critical for the final density profile, challenging current theory which emphasises compensation between the rates and durations of cell enlargement and wall thickening. These findings have implications for our understanding of how growth responds to environmental variability and the interpretation of tree rings as proxies of past climates. In addition, they provide a framework for the incorporation of explicit growth processes into models, such as those used to predict the role of vegetation in the future global carbon cycle. Finally, divergent responses in volume and mass with increasing temperature suggest caution in interpreting observations based on volume alone.

## Main

Lignified cellulose, “wood”, is the largest store of carbon in terrestrial plants, and as such plays a major role in the global carbon cycle^1^. Understanding controls on its formation is a major research priority^4,5^. Unlike most metabolic pathways, the process of wood formation, xylogenesis, is a synthetic dead-end without the potential to recycle materials, and is therefore highly regulated to avoid wasteful allocation of photosynthate^6^. This regulation results in conserved anatomical patterns, with tree rings in conifers and ring-porous species exhibiting transitions from low to high density wood each year. Nevertheless, these rings still carry the imprint of environmental conditions, a feature exploited in climate reconstructions, and as such underpin our ability to place contemporary climate change in an historical perspective^2^. However, despite the importance of this imprint for understanding past climates, and the relevance of wood formation for carbon sequestration and tree physiology, our understanding of how tree-ring anatomy arises and controls on its variability at intra- and inter-annual timescales remains poor.

Insights derived from studies on wood formation at the intra-annual timescale^7,8,9^, advances in understanding of cell size and growth regulation in meristems^10^, and high-resolution characterisation of biochemical and anatomical properties during wood development^11^, are combined here in a theoretical framework to analyse and explain how wood formation processes result in final ring structures. The framework explicitly considers xylogenesis in time and space, and thereby provides a mechanistic basis for understanding and predicting how the rate at which carbon is sequestered into wood varies with environmental conditions at daily to inter-annual timescales.

*Pinus sylvestris* L. (Scots pine) is used as the model species due to the good availability of critical observations. As in most conifers, its growth rings normally consists of earlywood with large cells and thin secondary walls, which transitions to latewood with small cells and thick walls in the final third or so of the ring. The total annual carbon increment is the result of integrating this density distribution over the total ring width, and cell lumen areas and wall thicknesses determine wood properties such as conductivity to water and strength. However, the mechanisms that create this pattern are not understood^9^. Recent observations of intra-annual cellular kinetics have quantified the relative contributions of different processes, such as cellular enlargement^7,12,13,9^, but have not yet been integrated into a general mechanistic explanation of how the final structures are created. A key question is why cellular density is relatively unresponsive to temperature, except in the final part of the developing ring. A paradigm that has emerged from statistical interrogation of intra-annual observations is that the lack of sensitivity in earlywood is due to compensation mechanisms between rates and durations, which somehow break down in latewood^14^.

The framework explored here combines observations, knowledge, and hypotheses for the fundamental mechanisms that underlie xylogenesis. The main assumptions are: (i) when not dormant, a single bifacial initial (stem) cell in each radial file grows radially and undergoes periclinal divisions, producing phloem outwards and xylem inwards^15^; (ii) these divisions are stochastically assigned to xylem or phloem, with prescribed probabilities^16^; (iii) when not dormant, inward cells grow radially, and can themselves divide if in the proliferation zone, with division-size regulation intermediate between a constant size and constant increment^10^ (this assumes that the vascular cambium and the apical meristem share regulatory mechanisms^17^); (iv) division is asymmetrical, with the relative growth rates of the daughters conditioned on their relative sizes, maintaining homeostasis^10^; (v) initial cells and their derivatives elongate in the radial direction at a rate dependent on temperature (calibrated to^18^; other factors such as turgor and nutrients are not considered here, but could be incorporated as controls on radial growth); (vi) once cells are no longer in the proliferation zone they continue to enlarge at a temperature-dependent rate while in an enlargement zone (calibrated to^11^); (vii) once they are no longer in the enlargement zone, secondary wall thickening occurs until they are no longer in the wall thickening zone, at which point they are assumed to lose their protoplasm and become mature, functioning, xylem; (viii) the rates of primary and secondary wall growth depend on temperature (calibrated to^18^), and the local availability of carbohydrates (calibrated to^11^); (ix) carbohydrates diffuse from the phloem along the radial file, with a prescribed phloem concentration and constant resistance between cells, creating an equilibrium profile between supply and demand each day (calibrated to^11^); (x) cells receive time-evolving positional information based on daylength that determines whether they are in the proliferation, enlargement, or wall thickening developmental zones (calibrated to^11^); (xi) the proliferation zone enters dormancy at a prescribed daylength^19^, and exits dormancy when a chilling day-conditioned heat sum is reached^20^ (calibrated to^11^).

### Overall density profile

The simulated distribution of cell-mass density across the growth ring at the end of the growing season compares favourably to observations (Fig. 1a), suggesting that its assumptions are highly plausible. No model parameter or underlying assumption was calibrated to observations of density distribution - the predicted pattern and absolute values are entirely an emergent property. In both observations and simulation, density remains low and fairly constant at c.0.4 g cm^−3^ for the first 40-50% of the ring, and then increases to a peak of c.0.85-1 g cm^−3^ in the final 10-30% of the ring, before declining in the final 10-20%.

**Figure 1.**
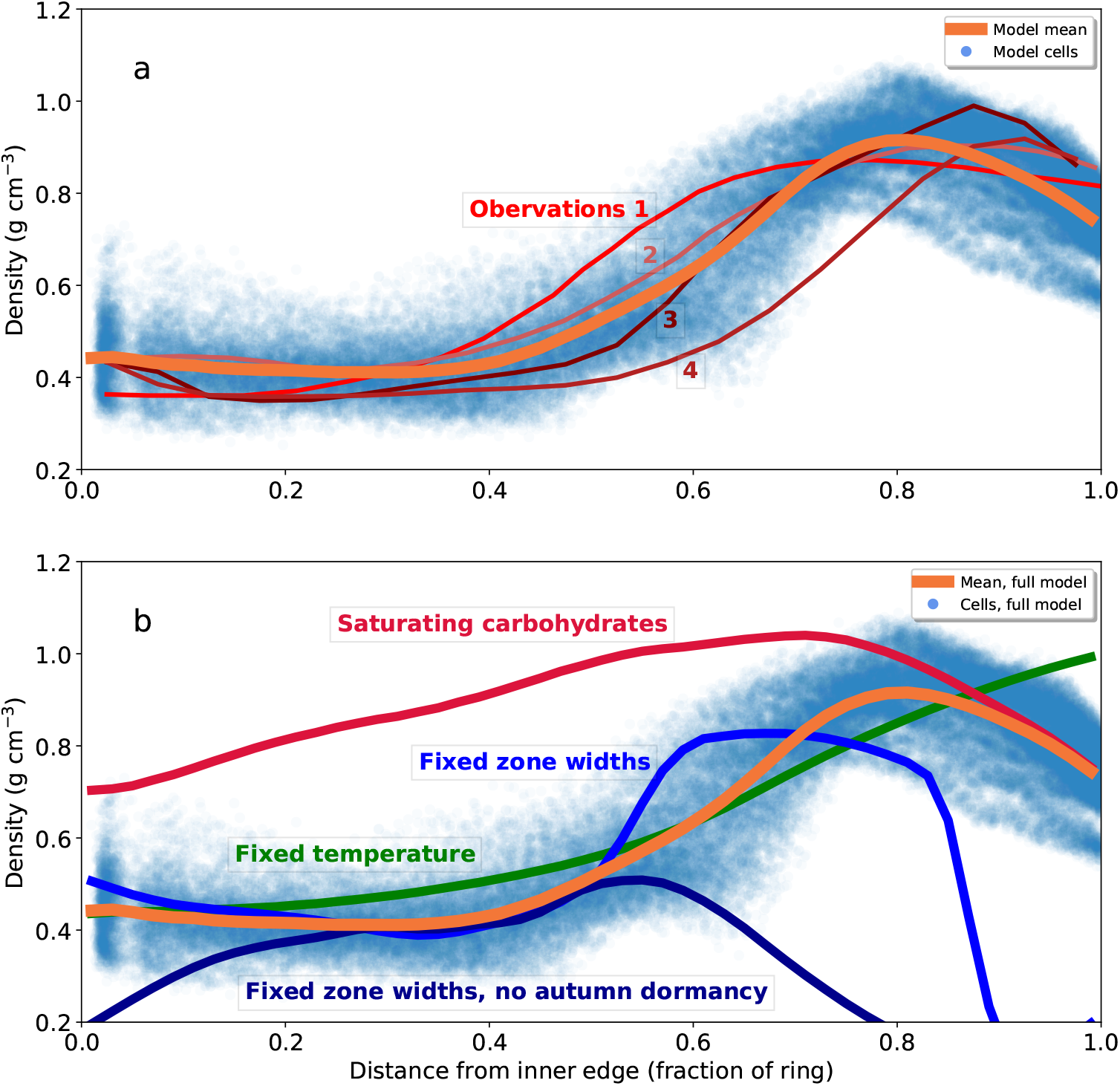
Simulated cell-mass densities as functions of relative distance across annual rings in *Pinus sylvestris* growing at 64.25 °N, 19.75 °E (northern Sweden), 1976-1995 using the full model. 100 radial files were simulated, with a spin-up over 1951-1975, and proliferating cells carried over between years. Symbols are individual cells and orange lines are 0.06-fraction running mean over 0.02-fraction bins. **a,** Comparison with observations of the same species from: 1. France^7^, morphometric density; 2. France^7^, X-ray; 3. southern Finland^21^, X-ray; 4. northern Finland^21^, X-ray. **b,** Effects of changing model assumptions on density. The fixed-temperature response uses 10 °C for cell enlargement and wall thickening; the fixed zone-widths responses keeps developmental zones at their values on 20 June, with or without autumn dormancy; and the saturating carbohydrates response assumes Δ*M* = Δ*M_max_* (Eqn. 8).

The mechanisms responsible for the observed intra-annual density pattern are not known, but could involve, singly or in combination, climate variability (e.g. temperature and/or water availability directly affecting cellular development), changing developmental zone widths/activity (possibly related to daylength signalling), and/or carbon supply (possibly related to variability in photosynthesis and whole-plant source/sink dynamics). Removing the effect of temperature on cell enlargement and wall thickening does not change the overall pattern of low-density earlywood transitioning to high-density latewood, but does smooth the rate at which density increases towards the latewood, and removes the density decline in the final 20% of the ring (Fig. 1b). Therefore although temperature seasonality is not responsible for the overall density pattern, some features, such as the existence of a peak in the latewood, are temperature-controlled. Removing the effect of temperature on wall thickening only also removes the late-season peak, demonstrating that it is caused by low late-season temperatures reducing the rate of wall thickening.

The effect of changing zone widths was investigated by fixing all widths at their values on 20 June for the whole year (Fig. 1b). The resulting profile still has low-density earlywood and an increase in density in latewood, suggesting that zone width variability is not the cause of the early-latewood transition. However, in contrast to the full model, the transition is steeper and there is a rapid decline in density at c.85% of the ring. Analysis of cellular dynamics through the year (cf. Figs. S1; S4) shows that this decline is due to cells stuck in the enlargement zone when it becomes dormant, and hence do not undergo any wall thickening. Removing autumn dormancy but keeping the zone widths fixed produces very wide rings because many additional cells are produced and enlarge, but there is no longer any high-density latewood (Fig. 1b). Therefore although zone width variability *per se* is not the cause of the earlywood-latewood transition, the cessation of enlargement due to late-season dormancy is required, increasing the duration of wall thickening relative to enlarging.

The influence of carbohydrates on the density profile was investigated by allowing wall thickening to continue at its maximum, temperature-limited, rate (Fig. 1b). While there is still an increase in density across most of the profile, and a peak in the latewood, earlywood has a much higher density than in the full model and the earlywood-latewood transition is much smoother. The gradual increase in density across the earlywood with saturating carbohydrates is due to the narrowing of the enlargement zone, reducing cell size (density is constant across the first 50% of the ring if zone widths do not change but still increases if temperature is fixed). The low density in earlywood cells in the full model is therefore due to their low carbohydrate concentrations during wall thickening as a result of their relatively large distance from the phloem (cf. Fig. S1). The role of compensation mechanisms in maintaining constant density constant across the earlywood is examined below. The densities across the latewood when carbohydrates are saturating and in the full mode converge towards the end of the ring. Late-season cells undergo wall thickening closer to the phloem than cells maturing earlier, resulting in lower carbohydrate limitation.

Taken together, these simulations demonstrate that the overall density profile in the ring, and hence its final carbon content, is determined by the spatial distribution of carbohydrates across the radial file, the contraction of the enlargement zone, and the imposition of autumn dormancy on the proliferation zone. The spatial distribution of carbohydrates is an emergent property of the developing ring, determined by the activities of the proliferation, enlargement, and wall thickening zones (Fig. S4).

### Temperature and cell-mass density

The model was used to investigate mechanisms for the observed high sensitivity of cell-mass density to temperature within the latewood^14^. Adding a fixed temperature increment of 2 K to all daily temperatures (but keeping dormancy forcing unchanged) results in a wider radial file, with the increase due to more earlywood (Fig. 2), as is observed with respect to the relationship between ring widths and the fraction of earlywood in diverse tree species^22,23^. Earlywood density is reduced with increased temperature, whereas latewood density is increased, consistent with observations^24^. Applying the temperature increment to wall thickening but not enlargement results in no change in earlywood density (Fig. 2), showing that the reduction in earlywood density is due to its effect on cell elongation, leading to cell size increasing proportionally more than mass. However, latewood density is increased to the same extent as when temperature is applied to both thickening and enlarging, showing that the latewood temperature response is due to the effect on thickening alone. The control on earlywood density from cell enlargement and latewood density from cell wall thickening is consistent with anatomical analysis in different species across a broad range of locations^24^.

**Figure 2.**
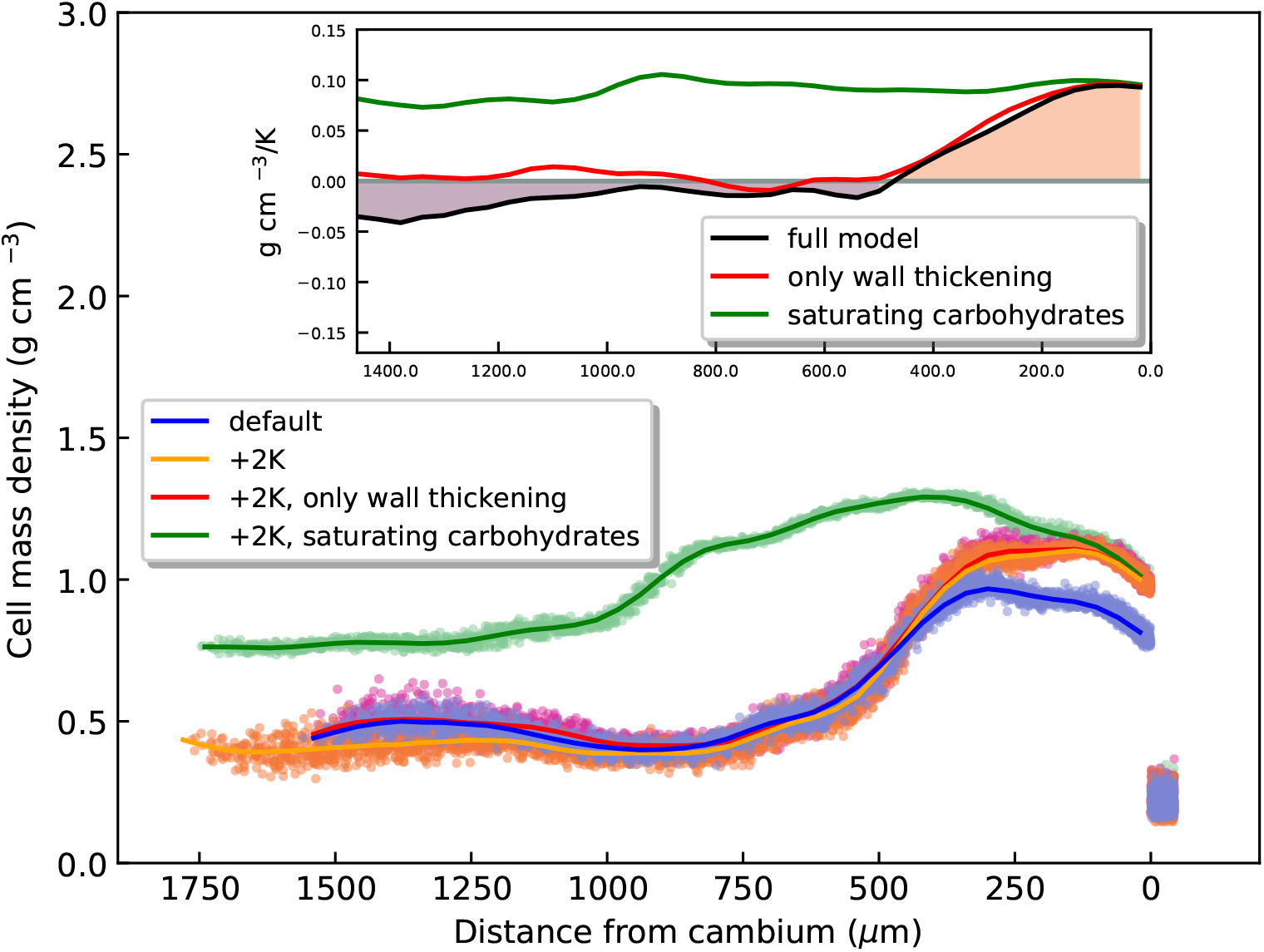
Effect of temperature on density across the simulated radial profile under different model configurations and forcings in *Pinus sylvestris* growing at the same site as in Fig. 1, 1995. Each point is an individual cell and solid lines are 120 *μ*m running means over 40 *μ*m bins. 100 radial files simulated for each model configuration and forcing. Temperature increment of +2 K applied to cell growth and wall thickening, or only wall thickening. Also shown is the result of applying the temperature increment to both cell growth and thickening but with the carbohydrate limitation to cell-wall growth removed. Inset shows profiles of ratio of change in density to change in temperature computed from the results in the main panel for the full model, only temperature effect on wall thickening, and with saturating carbohydrate and temperature effect on both enlargement and wall thickening.

The sensitivity of density to temperature, when applied to both enlarging and wall thickening, is negative in the earlywood but increases strongly through the latewood (inset in Fig. 2). This is consistent with the paradigm of compensation mechanisms occurring in the earlywood that eliminate the effects of temperature on density, with breakage of this compensation in the latewood^14^. The compensation mechanism has remained elusive, but has been interpreted as a consequence of tight negative coupling between rates and durations during xylogenesis^9^. However, if carbohydrates are assumed non-limiting, a different mechanism emerges. The relative effect of temperature on density under these conditions is nearly constant across the radial file, and equal to the maximum effect in the latewood for the full model (Fig. 2, inset). The low to negative effect of temperature in the earlywood of the full model is therefore attributable not to compensation between rate and duration, but rather to the limited supply of carbohydrates, similarly to how carbohydrate supply controls absolute density (Fig. 1). In contrast, high concentrations of carbohydrates in the latewood, due to its proximity to the phloem during wall thickening (Fig. S1), mean that wall building is relatively more limited by the temperature kinetics of the relevant biosynthetic machinery (Eqn. 9) than the substrate concentration (Eqn. 8). The cellular and zone-width dynamics responsible for the final ring anatomy are show in Fig. S1. Earlywood cells spend much less time in the wall thickening zone due to the high rate of proliferation and large width of the enlargement zone, quickly pushing the phloem outwards as the radial file lengthens. As the enlargement zone narrows over the growing season, cells spend more and more time in the wall thickening zone and, crucially, remain closer to the phloem. The cell walls of the latewood cells therefore become imprinted with temperature signals, leading to their utility as a climate proxy.

### Temperature and cell size

Higher temperatures lead to greater rates of cell enlargement and proliferation. The relative roles of these two processes in determining final cell sizes were quantified by comparing simulations in which temperature changes were applied to either the proliferation or enlargement zones, or to both.

Surprisingly, temperature increase results in a reduction in cell size across much of the ring (Fig. 3), demonstrating the existence of strong compensation mechanisms. The simulations in which temperature only influences the proliferation or enlargement zones shows that the compensatory mechanism is a subtle one. Higher temperatures in the enlargement zone indeed increase cell size, but higher temperatures in the proliferation zone decrease it. Higher rates of proliferation with increased temperature increase the rate at which cells are added to the enlargement zone (an average over the growing season of c.3 additional cells/file, +8%, for a 2 K rise), causing the duration of enlarging to decline, rapidly pushing the zones outwards (this can be envisaged in Fig. S1 as a steepening of the lines joining cells across the enlargement zone), compensating for the increased rate of enlargement.

**Figure 3.**
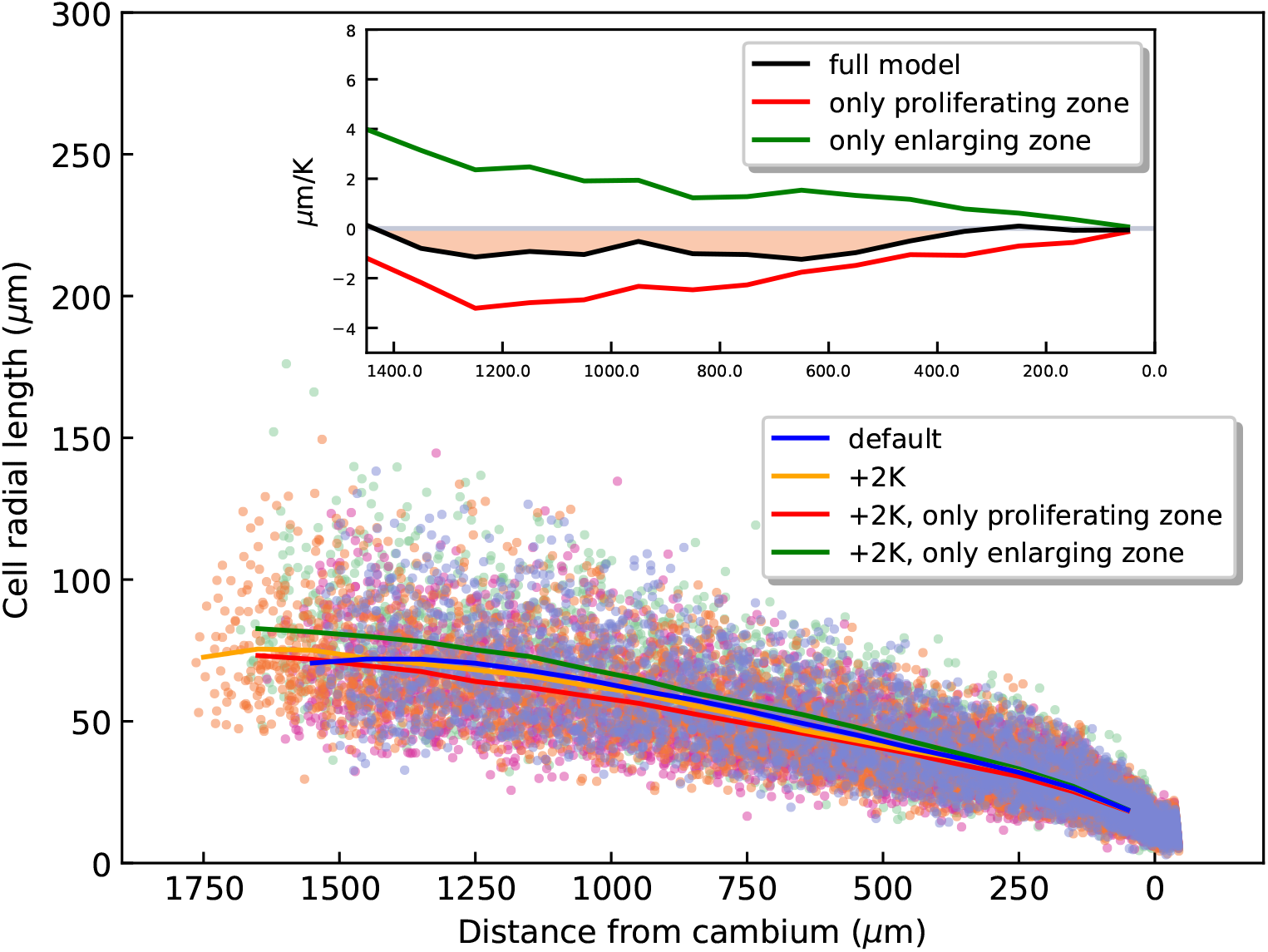
Effect of temperature on cell radial length across the radial profile under different model configurations and forcings in *Pinus sylvestris* growing at the same site as in Fig. 1, 1995. Each point is an individual cell and solid lines are 300 *μ*m running means over 100 *μ*m bins. 100 radial files were simulated for each model configuration and forcing. Temperature increment of +2 K applied daily to all cells, only proliferating cells, or only enlargement-zone cells. Inset shows radial profile of ratio of change in cell length to change in temperature over 0-2 K for the full model, only temperature effect on proliferating cells, and only temperature effect on enlargement-zone cells.

### Overall effects of temperature

A series of simulations was performed as above, but with step changes in temperature applied to the enlargement and wall thickening rates (Fig. 4). Ring width increases strongly and linearly with temperature across the entire range sampled. In contrast, total mass increases more strongly than ring width at negative temperature anomalies, but for higher temperatures it saturates, not increasing >+5 K. As was shown in Fig. 2, the increase in ring width with temperature is due to an increase in the width of earlywood, which causes overall density to decrease for positive temperature increments (Fig. 4; density decreases have been observed in different species over recent decades^23^). The increase in ring width is mainly due to an increase in the number of cells, rather than their size, which is consistent with the findings above of opposing effects of proliferation and enlargement rates on final cell sizes (Fig. 3). Removing the carbohydrate constraint on wall thickening causes total mass to increase very strongly with temperature as earlywood wall thickening is no longer limited by its distance from the phloem (this constraint increases with temperature due to the greater rate of increase in the cell-phloem distance; cf. Fig. S1). Maximum density increases at the same rate as total width at negative temperature anomalies, but gradually saturates as temperature increases.

**Figure 4.**
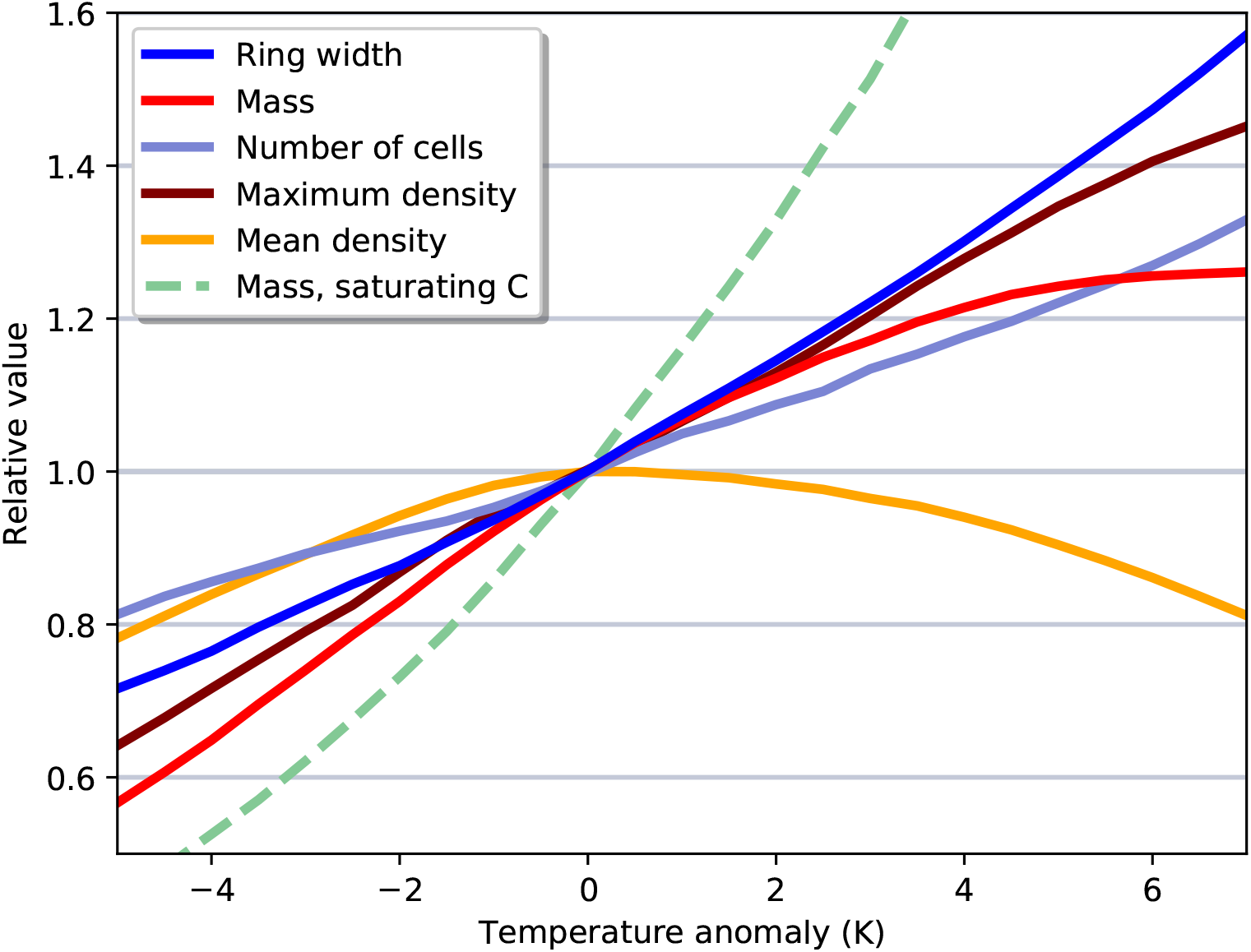
Temperature effects on simulated growth in *Pinus sylvestris* at the site used in Fig. 1, 1995. Each line is the 1.5 K running mean over 0.5 K increments for 100 radial files. Temperature anomaly applied as a step increment to the daily values used to compute cell enlargement and wall thickening.

The diverging responses of width and mass to temperature have significant implications for the measurement of tree mass growth and hence carbon uptake, which is usually inferred from volume increments^1^. In addition, the saturation of maximum density with temperature has implications for its use as a proxy for past climate, and may well play a role in the “divergence problem”^25^.

### Conclusions

The mechanisms explored here suggest that conifer ring structure results from the interactions of the activities and spatial configurations of developmental zones with the distribution of carbohydrates across the radial file. These interacting factors buffer the response of anatomical features to temperature variation, such as cell sizes, the earlywood-latewood transition, and earlywood density. Crucially, the diffusion of carbohydrates from the phloem plays a major role in the distribution of mass across the mature tree ring and its response to temperature. These findings have implications for our understanding of the response of carbon sequestration in trees to changing temperature and atmospheric CO_2_. Moreover, the framework presented here provides a mechanistic basis for the explicit treatment of tree growth in global vegetation models, enabling a balanced source/sink approach as advocated in a growing body of literature^26,27,5^. Such reformulation has the potential to reconcile inconsistencies between current models and observations (e.g. in relation to increasing CO_2_), with significant consequences for predictions of tree carbon storage, and hence the global carbon cycle.

## Methods

### Overall framework

Cells are arranged along radial files, with each cell in one of either the proliferation, enlargement, wall thickening, or mature zone, depending on its distance from the phloem. Only cells that contribute to the formation of xylem tracheids are treated. A daily timestep is used, on which cells in the proliferation and enlargement zones can enlarge in the radial direction if the zones are non-dormant, and on which secondary wall thickening occurs in the wall thickening zone. Cells in the proliferation zone divide if they reach a threshold radial length. Cell size control at division is intermediate between a critical size and a critical increment^10^. Mother cells divide asymmetrically, with the subsequent daughter relative growth rates inversely proportional to their relative sizes. Size at division and asymmetry of division are computed with added statistical noise^10^, and therefore the model is run for an ensemble of independent radial files with different initial conditions.

### Equations and parameters

#### Cell enlargement and division

Cells in the proliferation and enlargement zones, when not dormant, enlarge in the radial direction at a rate dependent on temperature and birth size relative to their sister. A Boltzmann–Arrhenius approach for the temperature dependence is used^28^:

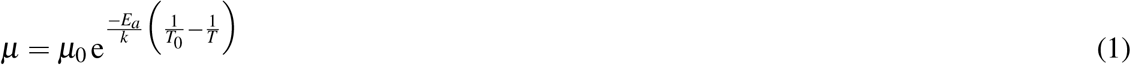

where *μ* is the relative rate of radial cell growth (*μ*m *μ*m^−1^ day^−1^), *μ*_0_ is *μ* at temperature *T_0_* (= 283.15 K), *E_a_* is the effective activation energy for cell enlargement, *k* is the Boltzmann constant (8.617×10^−5^ eV K^−1^), and *T* is temperature (K). *μ*_0_ was calibrated to the observed mean radial file length at the end of the elongation period^11^ (Table 1; see “Observations”), and *E_a_* was calibrated to the observed temperature dependence of annual ring widths^18^ (Table 1; Fig. S3; see “Observations”).

**Table 1.**
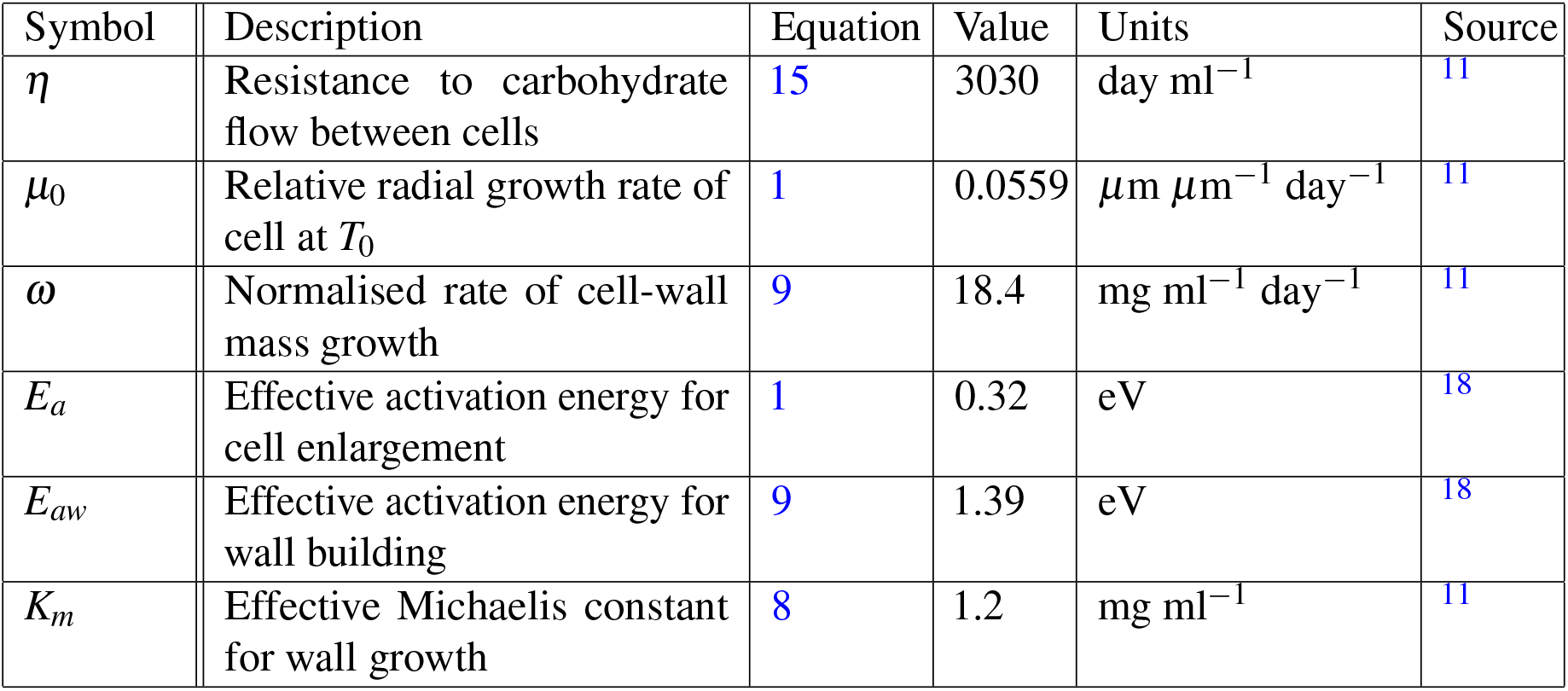
Parameters calibrated to observations. Calibration of *η*, *ω*, and *K_m_* uses predicted carbohydrate concentration at 257 *μ*m from the phloem, *μ*_0_ the final ring width, *E_a_* the ring width response to temperature, and *E_aw_* the maximum density response to temperature.

Radial length of an individual cell increases according to:

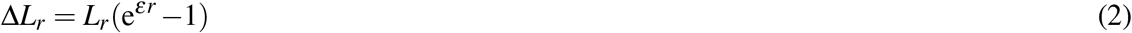

where Δ*L_r_* is the radial increment of the cell (*μ*m day^−1^), *L_r_* is the radial length of the cell (*μ*m), and ε is the cell’s growth dependence on relative birth size, given by^10^:

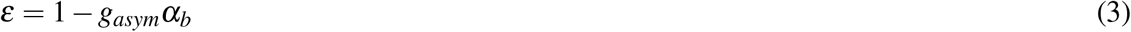

where *g_asym_* is the strength of the dependence of relative growth rate on asymmetric division (Table 2; unitless), and *α_b_* is the degree of asymmetry relative to the cell^10^ (scalar):

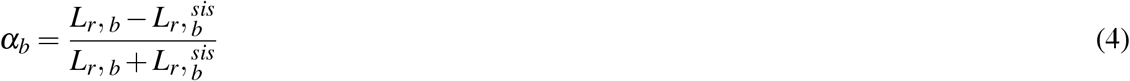

where *L_r, b_* is the radial length of the cell at birth ((*μ*m)) and 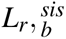 is the radial length of its sister at birth (*μ*m), which are calculated stochastically^10^:

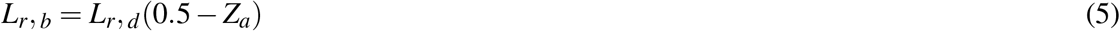

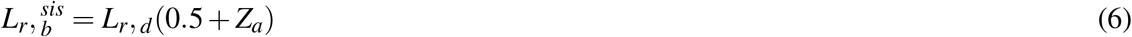

where *L_r, d_* is the length of the mother cell when it divides (*μ*m) and *Z_a_* is Gaussian noise with mean zero and standard deviation *σ_a_* (Table 2; −0.49 ≤ *Z_a_* ≤ 0.49 for numerical stability).

**Table 2.**
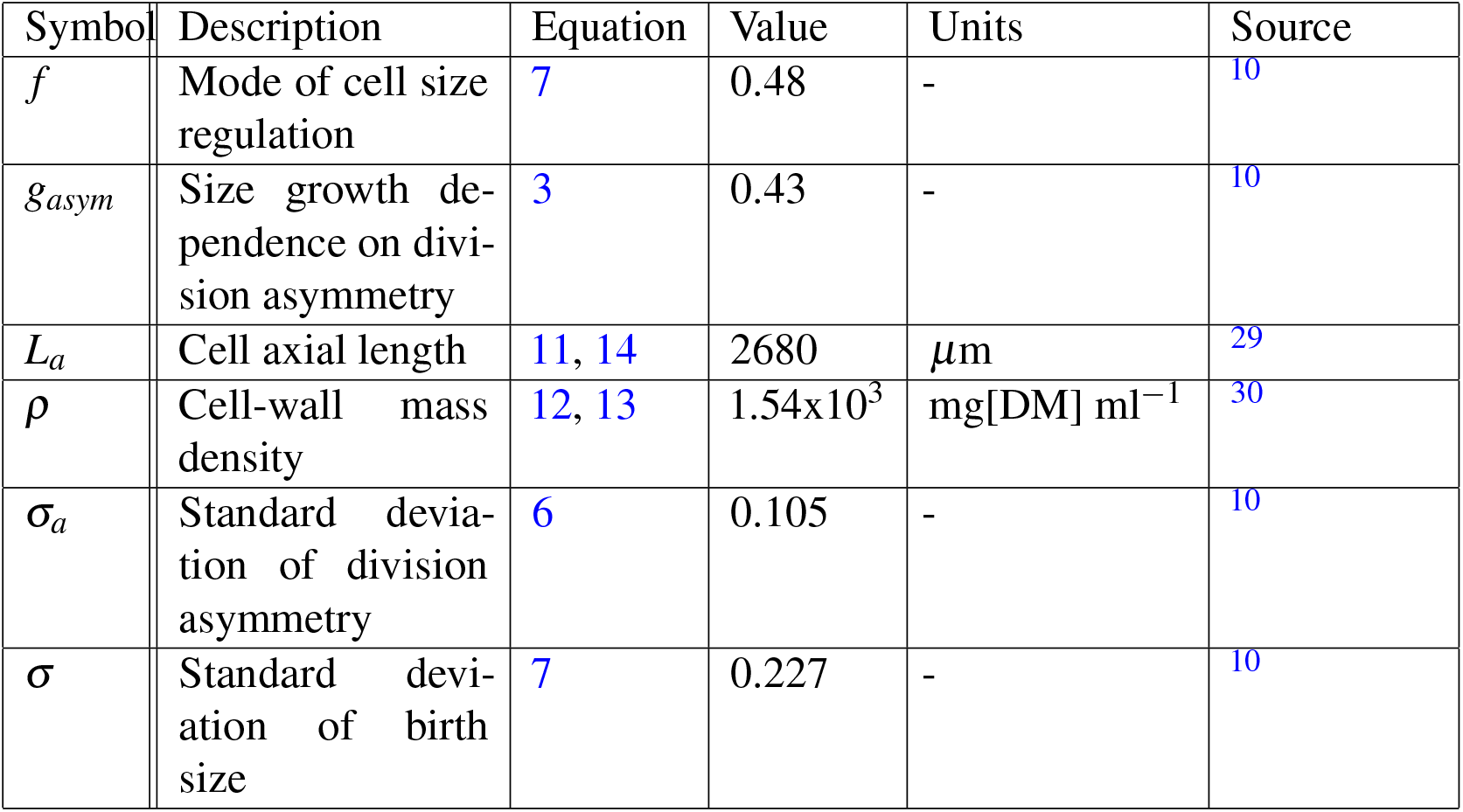
Parameters taken directly from literature.

Length at division is derived as^10^:

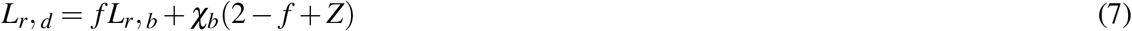

where *f* is the mode of cell-size regulation (Table 2; unitless), *χ_b_* is the mean cell birth size (Table 3; *μ*m), and Z is Gaussian noise with mean zero and standard deviation *σ* (Table 2).

**Table 3.**
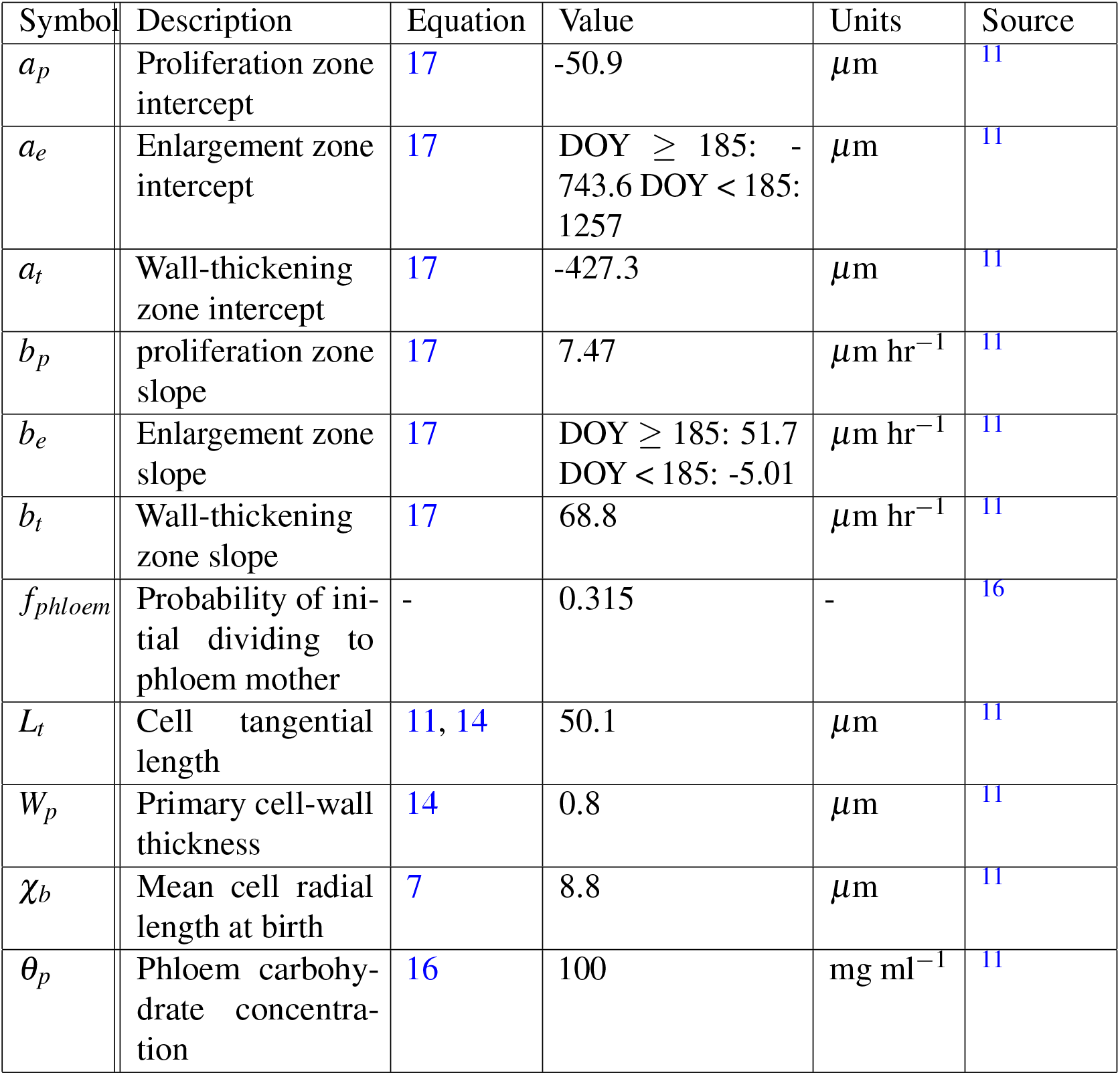
Parameters calculated from observations.

The first cell in each radial file is an initial, which produces phloem mother cells outwards and xylem mother cells inwards. It grows and divides as other cells in the proliferation zone, but on division one of the daughters is stochastically assigned to phloem or xylem, the other remaining as the initial. The probability of the daughter being on the phloem side is *f_phloem_* (Table 3).

#### Cell-wall growth

Both primary and secondary cell-wall growth are influenced by temperature, carbohydrate concentration, and lumen volume. A Michaelis-Menten equation is used to relate the rate of wall growth to the concentration of carbohydrates in the cytoplasm:

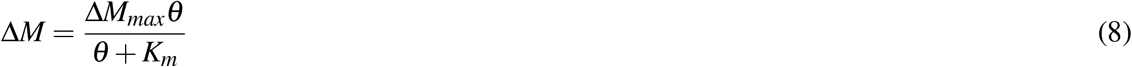

where Δ*M* is the rate of cell-wall growth (mg cell^−1^ day^−1^), Δ*M_max_* is the carbohydrate-saturated rate of wall growth (mg cell^−1^ day^−1^), *θ* is the concentration of carbohydrates in the cell’s cytoplasm (mg ml^−1^), and *K_m_* is the effective Michaelis constant (mg ml^−1^; Table 1).

The maximum rate of cell-wall growth is assumed to depend linearly on the lumen volume (a proxy for the amount of machinery for wall growth), and on temperature as in Eqn. 1:

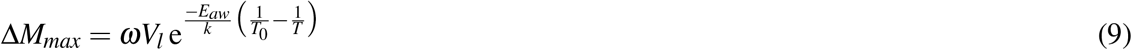

where *ω* is the normalised rate of cell-wall mass growth (i.e. the rate at *T*_0_; Table 1; mg ml^−1^ day^−1^), *V_l_* is the cell lumen volume (ml), and *E_aw_* is the effective activation energy for wall building (eV; Table 1; e). *ω* and *K_m_* were calibrated to the observed distribution of carbohydrates^11^ (see next section). *E_aw_* was calibrated to the observed temperature dependence of maximum density^18^ (Table 1; see “Observations”).

Lumen volume is given by:

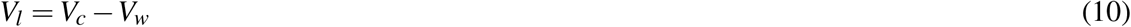

where *V_c_* is total cell volume (ml) and *V_w_* is total wall volume (ml). Cells are assumed cuboid and therefore *V_c_* is given by:

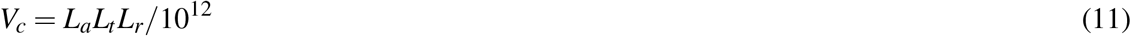

where *L_a_* is axial length (*μ*m; Table 2) and *L_t_* is tangential length (*μ*m; Table 3). *V_w_* is given by:

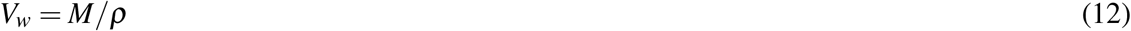

where *M* is wall mass (mg) and *ρ* is wall-mass density (Table 2; mg[DM]/ml).

Cells in the proliferation and enlargement zones only have primary cell walls. Δ*M_max_* (Eqn. 9) is therefore constrained to:

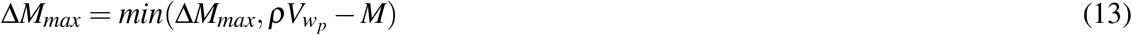

where *V_w_p__* is the required primary wall volume:

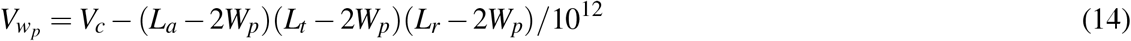

where *W_p_* is primary cell-wall thickness (Table 3; *μ*m).

#### Carbohydrate distribution

The distribution of carbohydrates across the radial file is calculated from the balance of diffusion from the phloem and the uptake into cell walls. The carbohydrate concentration in the phloem is prescribed, and the resulting concentration in the cytoplasm of the furthest living cell solved numerically. It is assumed that the rate of diffusion is fast relative to the rate of cell-wall building, and therefore concentrations are assumed to be in equilibrium. Carbohydrate diffusion between living cells is assumed to be driven by the concentration gradient:

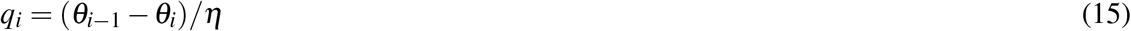

where *q_i_* is the rate of carbohydrate diffusion from cell *i* – 1 to cell *i* (mg day^−1^) and *η* is the resistance to flow between cells (calibrated to the observed distribution of carbohydrates^11^, see next section; Table 1; day ml^−1^).

At equilibrium, the flux into a given cell must equal the rate of wall growth in that cell plus wall growth in all cells further along the radial file. From this it can be shown that the equilibrium carbohydrate concentration in the furthest living cell in the radial file is given by:

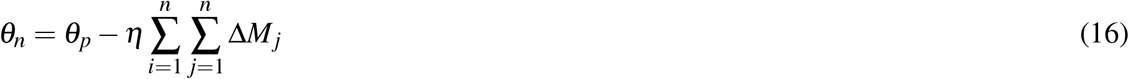

where *θ_p_* is the concentration of carbohydrates in the phloem (Table 3; mg ml^−1^) and *n* is the number of living cells in the file (phloem mother cells are ignored for simplicity). The rate of wall growth in each cell depends on the concentration of carbohydrates (Eqn. 8), and therefore *θ_n_* must be found that results in an equilibrium flow across the radial file. This is done using Brent’s method^31^ as implemented in the “ZBRENT” function^32^.

#### Zone widths

The widths of the proliferation, enlargement, and wall thickening zones vary in time, and are fitted to observations on three dates^11^ (see Fig. S1 and “Observations”). Linear responses to daylength were found, which are therefore used to determine widths for the observational period and later days:

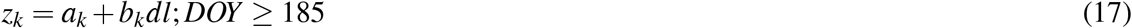

where *z_k_* is the distance of the inner edge of the zone from the phloem (*μ*m), *k* is proliferation (*p*), wall thickening (*t*), or enlargement (*e*), *a_k_* and *b_k_* are constants (Table 3), *dl* is daylength (s), and DOY is day-of-year. The proliferation zone width on earlier days when non-dormant was fixed at its DOY 185 width (assuming this to be its maximum, and that it would reach its maximum very soon after dormancy is broken in the spring). During dormancy, the proliferation zone width is fixed at its value on DOY 231 (the first day of dormancy^11^). The enlargement zone width prior to DOY 185 is assumed to be a linear extension of the rate of change after DOY 185. The wall-thickening zone width plays little role prior to DOY 185, and so was set to its Eqn. 17 value each earlier day. On all days the condition *z_t_* ≥ *z_e_* ≥ *z_p_* is imposed. Fig. S1 shows the resulting progression of zone widths through the year, together with the observed values.

#### Dormancy

Proliferation was observed to be finished by DOY 231^11^, and so the proliferation and enlargement zones are assumed to enter dormancy then. Release from dormancy in the spring is calculated using an empirical thermal time/chilling model^20^. It was necessary to adjust the asymptote and temperature threshold of the published model because the heat sum on the day of release calculated from observations in Sweden (see “Observations”) was much lower than reported for Sitka spruce buds in Britain in the original work:

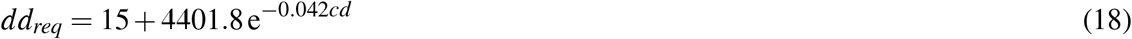

where *dd_req_* is the required sum of degree-days (°C) from DOY 32 for dormancy release and *cd* is the chill-day sum from DOY 306. The degree-day sum is the sum of daily mean temperatures above 0 °C, and the chill-day sum is the number of days with mean temperatures below 0 °C.

### Simulation protocols

Each simulation consisted of an ensemble of 100 independent radial files. Each radial file was initialised under dormant conditions by producing a file of cells with radial lengths *χ_b_* (1 + Z_a_), and then limiting these to only those falling in the proliferation zone on DOY 1. Values for ε (the relative growth of daughter cells) and *L_r, d_* (the radial length at division) were derived for each cell. The main simulations were conducted at the observation site in boreal Sweden (64.35 °N, 19.77 °E) over 1951-1995 to capture the growth period of the observed trees^11^. Results are mostly presented for 1995 when the observations were made. Simulations for calibration of the effective activation energies (i.e. *E_a_* and *E_aw_*) were performed at 68.26 °N, 19.63 °E in Arctic Sweden over 1901-2004^18^. Daily mean temperatures for both sites were obtained from the appropriate half-degree gridbox in a global dataset^33^.

### Observations

Observations of cellular characteristics and carbohydrate concentrations^11^ were used to derive a number of model parameters, and to test model output (model calibration and testing were performed using different outputs). Cell sizes, wall thicknesses, and positions in Figure 1^11^, an image of transverse sections on three sampling dates, were digitised using “WebPlotDigitizer“^34^. These, together with the numbers of cells in each zone and their sizes given in their text, were used to estimate zone widths, which were then regressed against daylength to give the parameters for Eqn. 17 (Table 3), mean cell size in the proliferation zone on the first sampling date (used to derive *χ_b_*; Table 3), mean cell tangential length (Table 3), and final ring width (used to calibrate *μ*_0_; Table 1). The thickness of the primary cell wall (Table 3) was derived by plotting cell wall thickness against time.

The distributions of carbohydrates along the radial files on the last sampling date for “Tree 1” and “Tree 3” (results for “Tree 2” are not shown for this date) shown in Figure 2^11^ were calculated. The masses for each of sucrose, glucose, and fructose in each 30 *μ*m section were digitised using the same method as for cell properties and then summed and converted to concentrations, with the results shown in Fig. S4. Mean observed carbohydrate concentration at a fixed point was used to calibrate values for the *η*, *ω*, and *K_m_* parameters in Table 1. Calibration was performed iteratively across the three parameters using graphical analysis of the relationships between their values and the predicted carbohydrate concentration at 257 *μ*m. Constraints were very tight for *η* and *ω*, but were poorer for *K_m_*.

The calibration target for the effective activation energy for wall deposition was the observed relationship between maximum density and temperature over 1901-2004 in northern Sweden^18^ (Fig. S2), and for the effective activation energy for cell enlargement the relationship between ring with and temperature in the same study (Fig. S3).

## Acknowledgements

This work has greatly benefited from discussions with numerous people, in particular Flurin Babst, Soumaya Belmecheri, Ulf Büntgen, Henri Cuny, Annemarie Eckes-Shephard, Patrick Fonti, David Frank, Andrew Hacket Pain, Christian Körner, Ben Poulter, and Valerie Trouet. The author acknowledges support from the Natural Environment Research Council under grant no. NE/P011462/1.

## Supplementary figures

**Figure S1.**
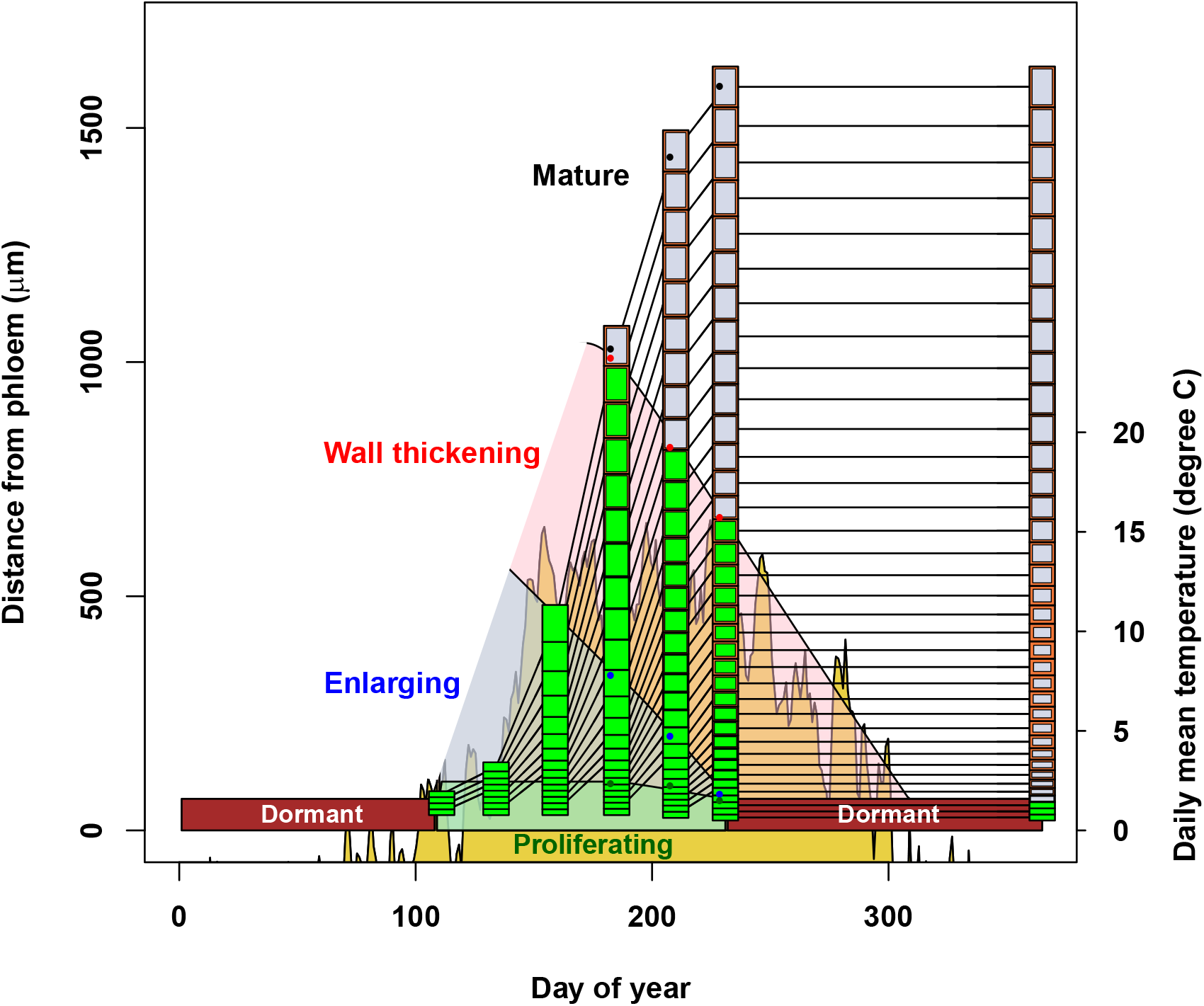
Simulated radial file development from the full-model simulation in Fig 1. Cells are means over 100 radial files. Cells positions are shown relative to the phloem, and are therefore represented as moving through development zones as they undergo differentiation. Filled circles are observed zone widths and total file length^11^ (see “Observations”). Dormancy affects both proliferation and enlarging cells. Living cells are shown with green protoplasm and mature, dead cells with grey lumen. Cell walls are orange and are drawn to the y-axis scale. The influence of temperature (yellow zone in background) on wall thickness is limited when the proliferation and enlargement zones are active due to the wall thickening zone rapidly retreating and the large distance from the phloem, reducing carbohydrate concentrations. In contrast, wall thickening in latewood cells occurs closer to the phloem and for longer, leading to higher densities and strong temperature effects.

**Figure S2.**
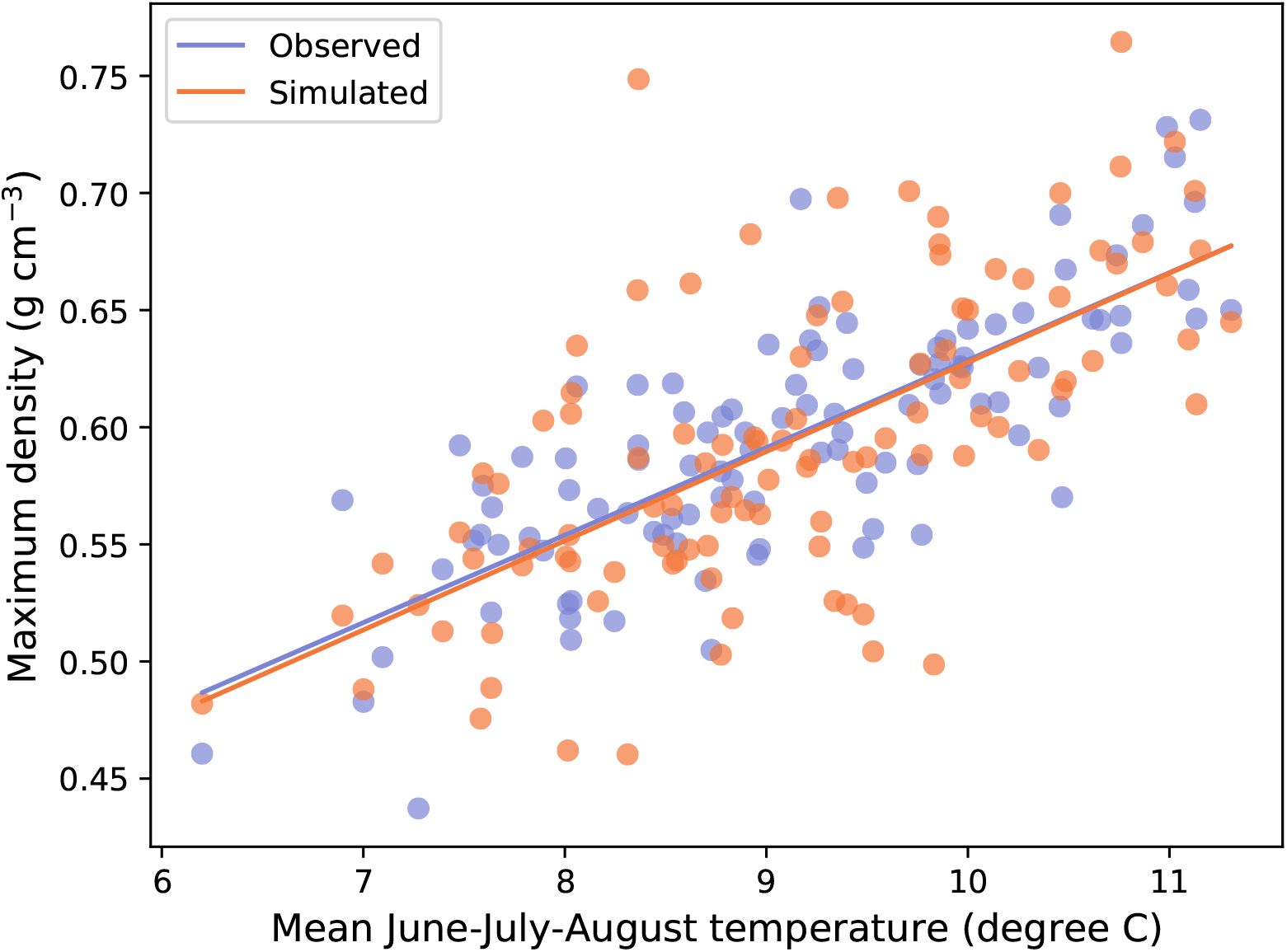
Observed and simulated relationships between annual maximum ring density and mean June-July-August temperature in northern Sweden. Simulation as for the full model in Fig. 1, except over 1901-2004, with latitude = 68.26 °N, and temperatures from the relevant half-degree grid-point in a global dataset^33^. Observations are “Tree Ring Maximum Density Chronology, Regional Curve Standardization” from ftp://ftp.ncdc.noaa.gov/pub/data/paleo/treering/reconstructions/europe/sweden/tornetrask-temperature2008.txt^18^, scaled using the observed mean of 0.615 g cm^−3^^18^. Model parameter *E_aw_* adjusted to achieve the same slope as observed (Table 1). Absolute densities were not calibrated.

**Figure S3.**
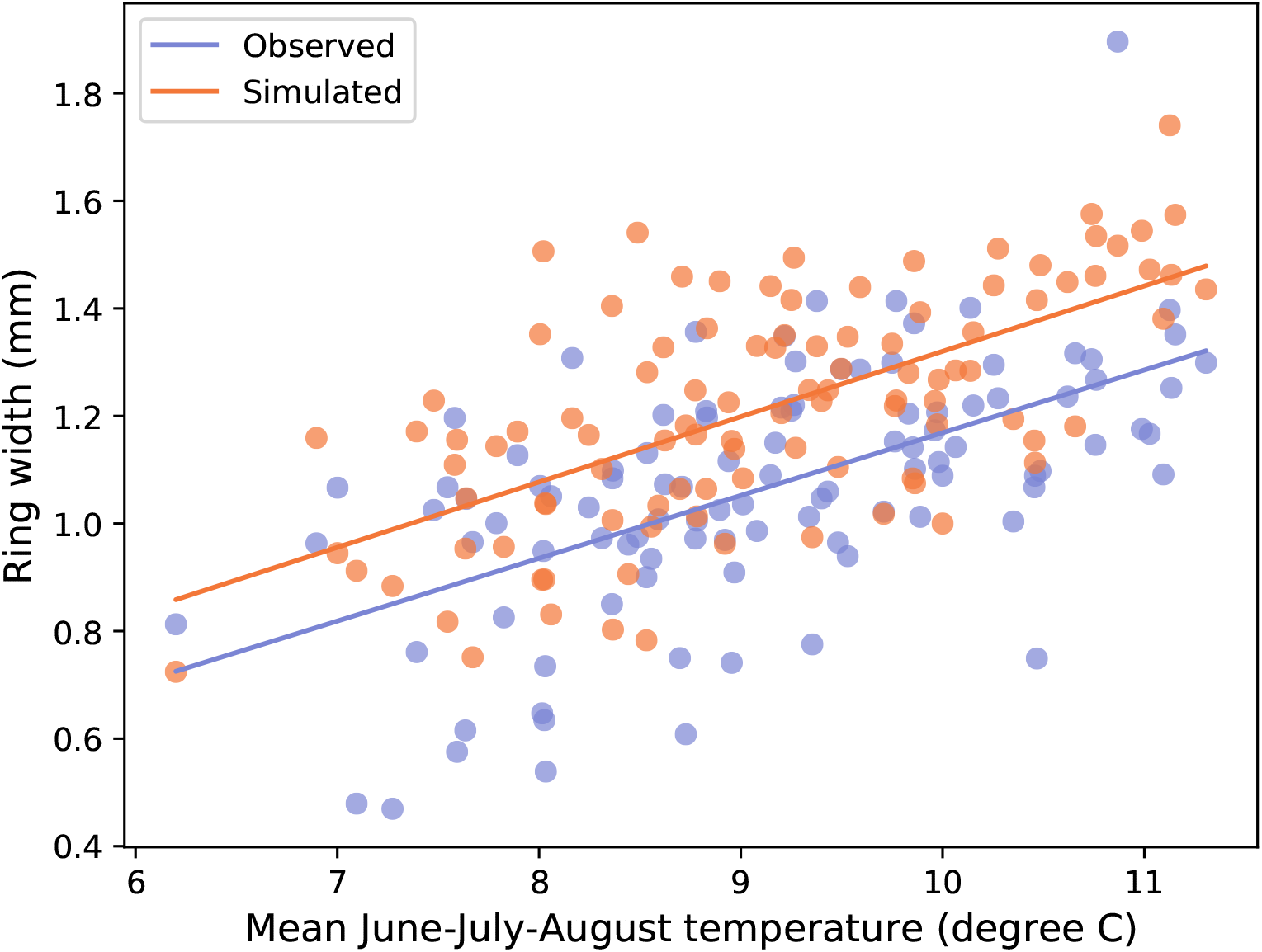
Observed and simulated relationships between annual ring width and mean June-July-August temperature in northern Sweden. Simulation undertaken as in Fig. S2. Observations are “Tree Ring Width Chronology, Regional Curve Standardization” from ftp://ftp.ncdc.noaa.gov/pub/data/paleo/treering/reconstructions/europe/sweden/tornetrask-temperature2008.txt^18^, scaled using the observed mean of 0.796 mm^18^. Model parameter *E_a_* adjusted to achieve the same slope as observed (Table 1). Absolute widths were not calibrated.

**Figure S4.**
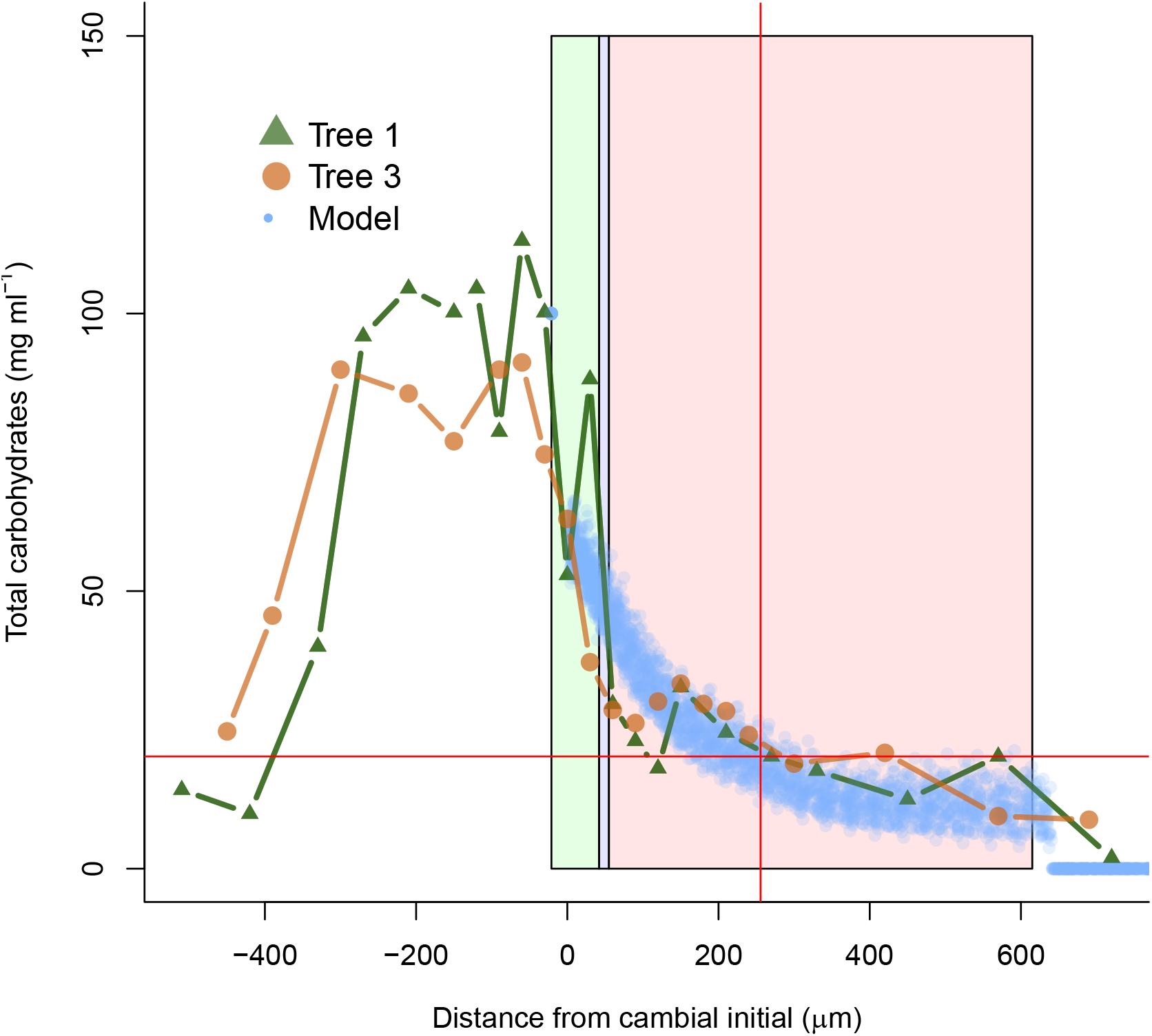
Observed^11^ and simulated distributions of carbohydrates across the developing wood on DOY 231. Model results are values for individual cells across 100 files, as for the full-model simulation in Fig. 1. Shaded light green area is the observed proliferation zone, shaded light blue area is the observed enlargement zone, and shaded light red area is the observed wall thickening zone. Horizontal and vertical red lines show the calibration target at 256 *μ*m.

## References

1. Pan, Y. et al. A large and persistent carbon sink in the world’s forests. Science 333, 988–993, DOI: 10.1126/science.1201609 (2011).

2. Mann, M. E., Bradley, R. S. & Hughes, M. K. Global-scale temperature patterns and climate forcing over the past six centuries. Nature 392, 779–787, DOI: 10.1038/33859 (1998).

3. Plomion, C., Leprovost, G. & Stokes, A. Wood gormation in trees. Plant Physiol. 127, 1513–1523, DOI: 10.1104/pp.010816 (2001).

4. Ursache, R., Nieminen, K. & Helariutta, Y. Genetic and hormonal regulation of cambial development. Physiol. Plantarum 147, 36–45, DOI: 10.1111/j.1399-3054.2012.01627.x (2013).

5. Friend, A. D. et al. On the need to consider wood formation processes in global vegetation models and a suggested approach. Annals For. Sci. 76, 49, DOI: 10.1007/s13595-019-0819-x (2019).

6. McCahill, I. W. & Hazen, S. P. Regulation of cell wall thickening by a medley of mechanisms. Trends Plant Sci. 24, 853–866, DOI: 10.1016/j.tplants.2019.05.012 (2019).

7. Cuny, H. E., Rathgeber, C. B. K., Frank, D., Fonti, P. & Fournier, M. Kinetics of tracheid development explain conifer tree-ring structure. New Phytol. 203, 1231–1241, DOI: 10.1111/nph.12871 (2014).

8. Cuny, H. E. et al. Woody biomass production lags stem-girth increase by over one month in coniferous forests. Nat. Plants 1, DOI: 10.1038/nplants.2015.160 (2015).

9. Cuny, H. E. et al. Couplings in cell differentiation kinetics mitigate air temperature influence on conifer wood anatomy. Plant, Cell & Environ. 42, 1222–1232, DOI: 10.1111/pce.13464 (2019).

10. Willis, L. et al. Cell size and growth regulation in the *Arabidopsis thaliana* apical stem cell niche. Proc. Natl. Acad. Sci. 113, E8238–E8246, DOI: 10.1073/pnas.1616768113 (2016).

11. Uggla, C., Magel, E., Moritz, T. & Sundberg, B. Function and dynamics of auxin and carbohydrates during earlywood/latewood transition in Scots pine. Plant Physiol. 125, 2029–2039, DOI: 10.1104/pp.125.4.2029 (2001).

12. Balducci, L. et al. Compensatory mechanisms mitigate the effect of warming and drought on wood formation: Wood formation under warming and drought. Plant, Cell & Environ. 39, 1338–1352, DOI: 10.1111/pce.12689 (2016).

13. Rathgeber, C. B. K., Cuny, H. E. & Fonti, P. Biological basis of tree-ring formation: a crash course. Front. Plant Sci. 7, DOI: 10.3389/fpls.2016.00734 (2016).

14. Cuny, H. E. & Rathgeber, C. B. Xylogenesis: coniferous trees of temperate forests are listening to the climate tale during the growing season but only remember the last words! Plant Physiol. 171, 306–317, DOI: 10.1104/pp.16.00037 (2016).

15. Shi, D., Lebovka, I., López-Salmerón, V., Sanchez, P. & Greb, T. Bifacial cambium stem cells generate xylem and phloem during radial plant growth. Development 146, dev171355, DOI: 10.1242/dev.171355 (2019).

16. Antonova, G. F. & Stasova, V. V. Seasonal development of phloem in scots pine stems. Russ. J. Dev. Biol. 37, 306–320, DOI: 10.1134/S1062360406050043 (2006).

17. Schrader, J. et al. A high-resolution transcript profile across the wood-forming meristem of poplar identifies potential regulators of cambial stem cell identity. The Plant Cell 16, 2278–2292, DOI: 10.1105/tpc.104.024190 (2004).

18. Grudd, H. Torneträsk tree-ring width and density ad 500–2004: a test of climatic sensitivity and a new 1500-year reconstruction of north Fennoscandian summers. Clim. Dyn. 31, 843–857, DOI: 10.1007/s00382-007-0358-2 (2008).

19. Petterle, A., Karlberg, A. & Bhalerao, R. P. Daylength mediated control of seasonal growth patterns in perennial trees. Curr. Opin. Plant Biol. 16, 301–306, DOI: 10.1016/j.pbi.2013.02.006 (2013).

20. Cannell, M. & Smith, R. Thermal time, chill days and prediction of budburst in *Picea sitchensis*. J. Appl. Ecol. 20, 951–963 (1983).

21. Decoux, V., Varcin, E. & Leban, J.-M. Relationships between the intra-ring wood density assessed by X-ray densitometry and optical anatomical measurements in conifers. Consequences for the cell wall apparent density determination. Annals For. Sci. 61, 251–262, DOI: 10.1051/forest:2004018 (2004).

22. Franceschini, T. et al. Decreasing trend and fluctuations in the mean ring density of Norway spruce through the twentieth century. Annals For. Sci. 67, 816–816, DOI: 10.1051/forest/2010055 (2010).

23. Pretzsch, H., Biber, P., Schütze, G., Kemmerer, J. & Uhl, E. Wood density reduced while wood volume growth accelerated in Central European forests since 1870. For. Ecol. Manag. 429, 589–616, DOI: 10.1016/j.foreco.2018.07.045 (2018).

24. Björklund, J. et al. Cell size and wall dimensions drive distinct variability of earlywood and latewood density in Northern Hemisphere conifers. New Phytol. 216, 728–740, DOI: 10.1111/nph.14639 (2017).

25. Briffa, K. R. et al. Reduced sensitivity of recent tree-growth to temperature at high northern latitudes. Nature 391, 678–682, DOI: 10.1038/35596 (1998).

26. Körner, C. Paradigm shift in plant growth control. Curr. Opin. Plant Biol. 25, 107–114, DOI: 10.1016/j.pbi.2015.05.003 (2015).

27. Fatichi, S., Pappas, C., Zscheischler, J. & Leuzinger, S. Modelling carbon sources and sinks in terrestrial vegetation. New Phytol. 221, 652–668, DOI: 10.1111/nph.15451 (2019).

28. Michaletz, S. T. Evaluating the kinetic basis of plant growth from organs to ecosystems. New Phytol. 219, 37–44, DOI: 10.1111/nph.15015 (2018).

29. Makinen, H. & Hynynen, J. Wood density and tracheid properties of Scots pine: responses to repeated fertilization and timing of the first commercial thinning. Forestry 87, 437–448, DOI: 10.1093/forestry/cpu004 (2014).

30. Dunlap, F. Density of wood substance and porosity of wood. J. Agric. Sci. II, 423–428 (1914).

31. Brent, R. P. Algorithms for minimization without derivatives. Prentice-Hall series in automatic computation (Prentice-Hall, Englewood Cliffs, N.J, 1972).

32. Press, W. H., Flannery, B. P., Teukolsky, S. A. & Vetterling, W. T. (eds.) Numerical recipes: the art of scientific computing (Cambridge University Press, Cambridge; New York, 1989).

33. Viovy, N. CRUNCEP Version 8 - Atmospheric Forcing Data for the Community Land Model. Research Data Archive at the National Center for Atmospheric Research, Computational and Information Systems Laboratory. https://doi.org/10.5065/PZ8F-F017. Accessed 11 July 2018. (2018).

34. Rohatgi, A. WebPlotDigitizer, v. 4.2. https://automeris.io/WebPlotDigitizer (2019).

